# Aberrant calcium signaling in astrocytes inhibits neuronal excitability in a human Down syndrome stem cell model

**DOI:** 10.1101/247585

**Authors:** Grace O. Mizuno, Yinxue Wang, Guilai Shi, Yizhi Wang, Junqing Sun, Stelios Papadopoulos, Gerard J. Broussard, Elizabeth K. Unger, Wenbin Deng, Jason Weick, Anita Bhattacharyya, Chao-Yin Chen, Guoqiang Yu, Loren L. Looger, Lin Tian

## Abstract

Down syndrome (DS) is a devastating genetic disorder causing severe cognitive impairment. The staggering array of effects associated with an extra copy of human chromosome 21 (HSA21) complicates mechanistic understanding of DS pathophysiology. We developed an in vitro system to examine the interplay of neurons and astrocytes in a fully recapitulated HSA21 trisomy model differentiated from DS patient-derived induced pluripotent stem cells (iPSCs). By combining calcium imaging with genetic approaches, we utilized this system to investigate the functional defects of DS astroglia and their effects on neuronal excitability. We found that, compared with control isogenic astroglia, DS astroglia exhibited more-frequent spontaneous calcium fluctuations, which reduced the excitability of co-cultured neurons. DS astrocytes exerted this effect on both DS and healthy neurons. Neuronal activity could be rescued by abolishing astrocytic spontaneous calcium activity either chemically by blocking adenosine-mediated astrocyte–neuron signaling or genetically by knockdown of inositol triphosphate (IP_3_) receptors or S100β, a calcium binding protein coded on HSA21. Our results suggest a novel mechanism by which DS alters the function of astrocytes, which subsequently disturbs neuronal excitability. Furthermore, our study establishes an all-optical neurophysiological platform for studying human neuron-astrocyte interactions associated with neurological disorders.

**Significant statement:** Down syndrome (DS) is the most common genetic disorder caused by trisomy of chromosome 21 (HSA21). Problems with cognitive impairment, have not been properly addressed due to the inability to fully recapitulate HSA21, which is further confounded by the snapshot views of morphological changes of brain cells in isolation obtained from current studies. The brain develops neural networks consisting of neurons and glial cells that work together. To understand how DS affects the neural networks, we used DS patient-derived stem cells and calcium imaging to investigate functional defects of DS astrocytes and their effects on neuronal excitability. Our study has significant implication in understanding functional defects during brain development underlying DS.

## Introduction

Down syndrome (DS) is a neurodevelopmental disorder occurring in 1 in 750 live births worldwide. DS is caused by trisomy of chromosome 21 (Ts21)^1^, leading to triplication of up to 400 genes, resulting in an array of phenotypes, including profoundly impaired cognitive function. The brains of DS patients demonstrate consistent pathophysiological changes, such as reduced volume, altered neuronal densities and structure, and disturbed balance of all cell types. Confronted with this genetic complexity, it is difficult to determine precise molecular and cellular mechanisms of disease establishment and maintenance. Consequently, there are no therapeutic approaches to mitigate the effects of DS.

To date, DS pathophysiology has been primarily studied in rodent models, (e.g. Ts65Dn, Ts1cje and Ts1Rhr)^2^. Though useful information has been revealed, rodent models do not faithfully reproduce DS pathophysiology, due in part to incomplete synteny between HSA21 and the homologous mouse regions. Furthermore, rodent modeling of complex neurodevelopmental disorders such as DS is limited by the fact that the human brain is far more complicated than the rodent brain in terms of structure of the neural circuitry, plasticity, and cognitive capacity.

Advances in induced pluripotent stem cell (iPSC) technology have enabled the modeling of complex diseases such as DS in the context of human cell biology^3,4^. These models are highly desirable for understanding disease neuropathophysiology and for developing therapeutics. By culturing iPSCs from DS individuals it is possible to achieve full expression of the human HSA21 region. In addition, the use of isogenic control lines eliminates inter-individual variability, restricting genotype differences solely to HSA21 dosage.

Recently, two Ts21-iPSC derived DS models have been reported. Weick et al. established Ts21-iPSC lines from two sets of human fibroblasts and differentiated them into neurons. They found that Ts21-neurons displayed reduced synaptic activity compared to control neurons, while maintaining the ratio of differentiated excitatory and inhibitory neurons. Chen et al., on the other hand, engineered Ts21 iPSCs from a different human fibroblast line and reported that conditioned medium from Ts21-iPSC derived astroglia had a toxic effect on neuronal maturation and survival. Although these two elegant studies provide complementary perspectives on the defects of human neurons or astroglia associated with DS, they studied neurons and astrocytes in isolation. Growing evidence suggests that astrocytes substantially contribute to neurological and psychiatric disorders by affecting neuronal function^5–9^. Indeed, astrocytes have been implicated in multiple rodent studies as playing an important role in DS^10,11^. A number of genes involved in DS, including TSP-1 and APP have been shown to be expressed in astrocytes and have been implicated in Alzheimer’s disease^12,13^. A complete mechanistic understanding of DS pathophysiology requires studying the communication between neurons and astrocytes at the network level.

Unlike neurons, whose excitable membranes allow action potentials to be transmitted cell-wide within milliseconds, astrocyte-wide signaling occurs *via* intracellular calcium (Ca^2+^) transients lasting for seconds^14^. These intracellular Ca^2+^ transients can be triggered by neuronal activity^15^ and are thought to induce release of gliotransmitters such as glutamate, GABA, ATP and D-serine^16–19^, which in turn modulate neural activity. Although gliotransmitter identity and release mechanisms are controversial^20–22^, intracellular Ca^2+^ dynamics are generally acknowledged to encode astrocyte activity. More importantly, altered astrocyte calcium dynamics were reported in cultured cells from the rodent DS models^13,23^.

Based on these previous studies, we hypothesized that DS could affect neuronal excitability through altered astrocytic Ca^2+^ dynamics, leading to alterations in astrocyte-neuron signaling pathways. Therefore, we differentiated the Ts21-iPSC lines reported in Weick et al. to astrocytes and neurons to establish a novel Ts21-iPSC-derived neuron-astrocyte co-culturing system to uncover functional deficits of neural networks. We focused on astrocytic Ca^2+^ dynamics and the specific interactions between astrocytes and neurons. We show that aberrant Ca^2+^ fluctuations in human DS astrocytes reduce the excitability of co-cultured human neurons and alter their synaptic properties. These effects are mediated by overexpression of the HSA21 protein S100β. Our study explores the contribution of astrocytes to abnormal neural circuit development, beyond the traditional view of trophic, supporting roles, and suggests causal roles of HSA21 gene overload in DS etiopathogenesis.

## Results

### Generation and differentiation of astroglia from human Ts21 iPSCs

Using established protocols^24^, we differentiated astroglia from previously reported iPSC lines by Weick et al., DS1 and DS4, which are trisomic for chromosome 21, and DS2U, a control isogenic line (**Supplementary Fig. 1a-b**)^4^. After 120 days, all three iPSC lines robustly expressed astrocyte precursor marker CD44, mature astrocyte markers glia fibrillary acidic protein (GFAP) and aquaporin 4 (AQP4), as determined by immunofluorescence and confirmed by quantitative reverse transcription PCR (qPCR) (**Supplementary Fig. 1d-f, Supplementary Table 1**). Karyotype analysis prior to and after experiments confirmed trisomy of DS1- and DS4-derived astroglia (DS1A and DS4A) and disomy of DS2U-induced astroglia (DS2UA) (**Supplementary Fig. 1b**). Using qPCR, we further observed global expression of a panel of astrocyte specific markers such as excitatory amino acid transporter 1 (*EAAT1*), aldolase C (*ALDOC*), connexin-43 (*CX43)*, *SOX9*, and nuclear factor I A (*NFIA*) in all three lines (**Supplementary Fig. 1c**)^9^, indicating successful astroglia differentiation of the iPSCs. Consistent with previous reports, DS astroglia showed increased expression levels of HSA21 genes compared to control astroglia, including *S100β*^25^, amyloid beta precursor protein (*APP)*^23^ and transcription factor *ETS2*^26^, as well as higher levels of non-HSA21 genes associated with oxidative stress, such as catalase (*CAT)*^27^ and quinone oxidoreductase (*CRYZ)*^4^(**Supplementary Fig. 1c**). Morphologically, DS astroglia occupied larger territories than DS2UA; the total arborization size of DS astrocytes was significantly greater than that of control isogenic astroglia (**Supplementary Fig. 1g**).

### DS astroglia inhibit the excitability of co-cultured neurons

We next studied the potential influence of DS astroglia on co-cultured neurons. Using established protocols^28,29^, three lines of cortical TUJ1^+^ (neural-specific β-III tubulin) neurons were differentiated from the DS1 and DS2U iPSC lines and a control H9 human embryonic stem cell (hESC) line (**Supplementary Fig. 2a–b**). Differentiated neurons were infected with lentivirus encoding GCaMP6m driven by the neuron specific promoter *synapsin-1* (**Supplementary Fig. 2c**). To establish a baseline of neuronal excitability, we monitored fluorescence changes in neurons in response to a series of electrically evoked field potentials (FPs) in the absence of astrocytes. The magnitude of evoked Ca^2+^ transients in neurons increased with the number of applied FPs (Fig. 1a). Evoked signals were abolished by addition of 1 μM tetrodotoxin (TTX; a voltage-gated sodium channel blocker) (Fig. 1a), suggesting that Ca^2+^ signals in neurons were triggered by action potentials. The expression of multiple voltage-gated sodium-channel isoforms in differentiated neurons was confirmed by qPCR assay (**Supplementary Fig. 2d**).

After confirming the basis of neuronal excitability, we recorded neuronal activity when co-cultured with DS1-, DS4-, or DS2U-derived astroglia, as well as human primary astrocytes (HA). H9 hESC-derived neurons co-cultured with DS astroglia (DS1A or DS4A) showed significantly decreased FP-evoked Ca^2+^ amplitudes relative to neurons cultured alone (normalized ΔF/F; DS1A: 0.63±0.06, *P*=0.0042; DS4A: 0.57±0.05, *P*<0.001), whereas neurons co-cultured with control isogenic astrocytes (DSU2A) or human primary astrocytes were not significantly affected (DS2UA: 1.00±0.04, *P*=0.93; HA: 0.88±0.04, *P*=0.059; Fig. 1b).

**Figure 1.**
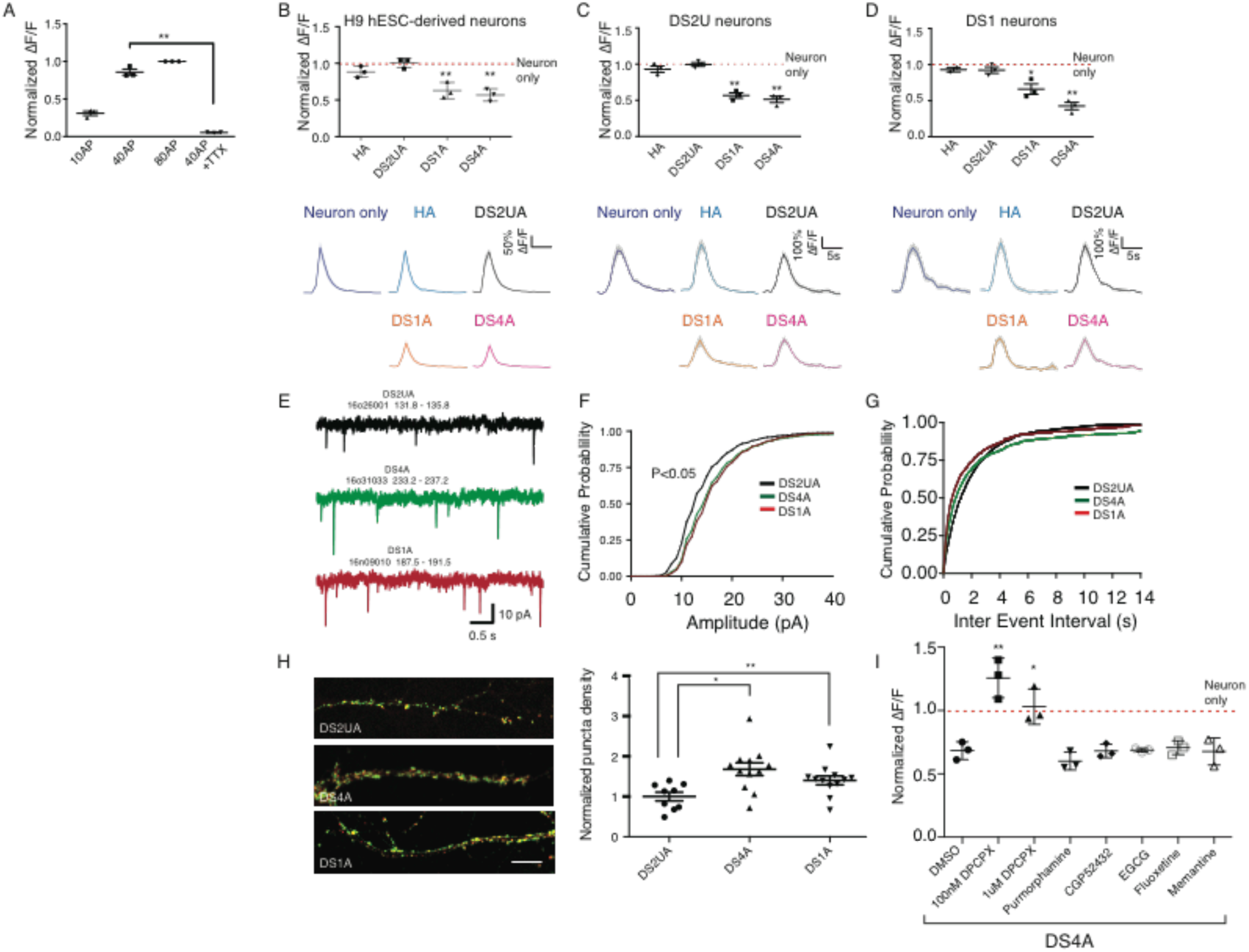
DS astroglia inhibit neuronal excitability during co-culture. (a) The fluorescence changes (ΔF/F) of H9 hESC-derived neurons in response to a variety of FP stimuli; ΔF/F at 10 FPs, 40 FPs, and 40 FPs in the presence of 1 μM TTX were normalized to ΔF/F at 80 FPs. (b-d) The responses of H9 hESC- (b), isogenic DS2U- (c), and DS1-iPSC- (d) derived neurons to FP stimuli (40 FPs at 30Hz) when co-cultured with or without astroglia. ΔF/F induced by FP stimuli in the presence of astrocytes was normalized to that of neurons alone (red dotted lines). Representative traces showing Ca^2+^ transients triggered by FPs in neurons are shown (right panel). (e) Example recordings of mEPSCs from 1 neuron from each group. (f) Cumulative probability of the mEPSC amplitude shifted rightward in both DS4A and DS1A groups compared with the DS2UA group. (g) No change was seen in the cumulative probability of the mEPSC inter-event interval. (h) Representative images and quantification of puncta density expressing both pre-synaptic protein synapsin and post-synaptic scaffolding protein PSD95 on oblique dendrites of co-cultured rat hippocampal neurons. (i) The fluorescence changes of H9 hESC-derived neurons in response to 40 FPs stimuli when co-cultured with DS4A in the presence of DMSO or a series of drugs are shown and normalized to changes when co-cultured with DS2UA. Several compounds showing therapeutic effect in DS mouse models had no rescuing effect on neuronal activity when co-cultured with DS4A, except DPCPX, an A1-receptor antagonist. The fluorescence changes of H9 hESC-derived neurons, in response to 40 FPs stimuli when co-cultured with DS4A in the presence of DMSO or a series of drugs, are shown and normalized to fluorescence changes when co-cultured with DS2UA.

Similar neuronal-activity suppression imposed by DS astroglia was also observed in neurons derived from the two other iPSC lines. DS2U derived neurons co-cultured with DS astroglia (DS1A or DS4A) showed significantly decreased FP-evoked Ca^2+^ amplitudes relative to neurons cultured alone (normalized ΔF/F; DS1A: 0.57±0.04, *P*<0.001; DS4A: 0.51±0.04, *P*<0.001; DS2UA: 0.99±0.03, *P*=0.89; HA: 0.93±0.04, *P*=0.18; Fig. 1c). Likewise, DS1 derived neurons co-cultured with DS astroglia (DS1A or DS4A) showed significantly decreased FP-evoked Ca^2+^ amplitudes relative to neurons cultured alone (normalized ΔF/F; DS1A: 0.66±0.07, *P*=0.0092; DS4A: 0.43±0.05, *P*<0.001; DS2UA: 0.92±0.04, *P*=0.15; HA: 0.95±0.03; *P*=0.18; Fig. 1d). Decreased neuronal activity in the presence of DS astroglia was observed under a variety of stimulation conditions, but was most prominent during modest stimulation such as 10FPs (**Supplementary Fig. 2e**). Taken together, DS astroglia inhibited neuronal excitability of neurons derived from either trisomy or disomy iPSC lines.

In addition, all co-cultured astrocytes significantly accelerated decay-to-baseline of evoked neuronal Ca^2+^ transients (T_0.5_=1.62±0.14 for neuron-alone; T_0.5_=1.22±0.08, 1.25±0.12, 1.11±0.13, and 1.18±0.1 for neurons co-cultured with HA, DS2UA, DS1A, and DS4A respectively, *P*<0.01; **Supplementary Fig. 2f**), presumably because astrocytic glutamate clearance following FP-evoked release occurs at similar rates.

### DS astroglia promote synaptic connectivity

As DS astroglia suppress neuronal activity, we next sought to determine if DS astroglia influence synaptic function. DS astroglia were co-cultured with dissociated rat hippocampal neurons, and miniature excitatory post-synaptic currents (mEPSCs) were recorded in the presence of TTX, NMDA receptor antagonist D-AP5, and GABA_A_ antagonist bicuculline, to isolate the fast AMPA receptor-mediated mEPSC component (Fig. 1e–g, **Supplementary Fig. 2g-h**). Cumulative distribution plots showed that the mean amplitude of mEPSCs was significantly larger in neurons co-cultured with either DS4A and DS1A groups compared with control DS2UA (DS2UA: 14.21±0.42; DS1A: 16.35±0.78, *P*=0.032; DS4A: 16.26±0.73, *P*=0.019; Fisher’s least-significant difference test) (Fig. 1f, **Supplementary Fig. 2g**, *P*<0.05). mEPSC frequency was similar in all three groups, with a trend towards higher mEPSC frequencies in the neurons co-cultured with DS4A and DS1A groups (*P*=0.204; DS2UA: 0.56±0.06; DS1A: 1.29±0.45; DS4A: 1.10±0.36) (Fig. 1g, **Supplementary Fig. 2h**).

We next evaluated the effects of human astroglia on synapse formation using quantitative image analysis^30^. We quantified the density of punctae expressing both the pre-synaptic protein synapsin-I and the post-synaptic scaffolding protein PSD95 on oblique dendrites of rat hippocampal neurons co-cultured with astroglia. We found that synapse density significantly increased by 1.5- and 1.3-fold in neurons co-cultured with DS astrocytes (DS1A, (*P*=0.0039); DS4A, (*P*=0.02), respectively) compared with those co-cultured with isogenic control astrocytes (Fig. 1h). Taken together, these results suggest that DS astroglia are capable of modulating neuronal excitability, as well as synaptic activity and density.

### Pharmacological rescue of suppressed neuronal excitability

We next examined whether pharmacological drugs that block astrocyte-neuron communication could rescue the suppressed neuronal excitability. We examined a panel of small molecule drugs that have been shown to rescue synaptic abnormalities in DS mouse models^31^. These compounds, including purmorphamine (sonic hedgehog agonist), CGP52432 (GABA_B_R antagonist), epigallocatechin-3-gallate (EGCG, DYRK1A inhibitor), fluoxetine (serotonin reuptake inhibitor), and memantine (NMDA receptor antagonist) failed to rescue decreased neuronal activity associated with DS astroglia (normalized ΔF/F=0.60±0.04, 0.68±0.03, 0.70±0.02, 0.71±0.03, and 0.68±0.06, from purmorphamine to memantine; *P*=0.22, 0.95, 0.73, 0.67, 0.93; *n*=3; Fig. 1i).

Next we examined the chemical transmitter ATP, since astrocytic release of ATP has been shown to modulate synaptic function, with intracellular Ca^2+^ transients increasing probability of release^16,18,19^. To what extent ATP potentiates and/or inhibits neuronal activity is still under debate; however, adenosine, a rapid ATP breakdown product, has been shown to inhibit synaptic activity via G_i_-coupled A_1_ adenosine receptors^32–36^. To test whether suppressed neuronal excitability is caused by adenosine-mediated signaling, we treated H9 neurons co-cultured with DS astroglia (DS4A) with an adenosine receptor antagonist, followed by imaging FP-evoked neuronal activity. In particular, the A_1_ receptor antagonist DPCPX fully rescued suppressed neuronal activity, especially at lower concentrations (100 nM: normalized ΔF/F=1.20±0.09, *P*=0.004; 1 μM: 0.98±0.08, *P*=0.018; *n*=3; Fig. 1i). This suggests that the suppressed neuronal excitability is influenced by purinergic signaling.

### DS astroglia exhibit abnormally frequent spontaneous Ca^2+^ fluctuations

Astrocytic Ca^2+^ signaling has been proposed to modulate neural-circuit activity and structure^37,38^; the suppressed excitability of neurons was specific to DS astroglia and could be rescued when astrocyte-neuron communication was blocked by an adenosine receptor antagonist. This evidence led us to further investigate calcium dynamics in astroglia. We focused on optical recordings of calcium dynamics in astroglia using the genetically encoded indicator GCaMP6m^39^. We used the machine-learning software Functional Astrocyte Phenotyping (FASP)^40^ to facilitate automated detection and analysis of complex Ca^2+^ dynamics in astroglia.

The differentiated astroglia indeed displayed prominent spontaneous Ca^2+^ transients, which were frequently periodic and especially apparent in DS astroglia (Fig. 2a, **Supplementary Movie 1&2**). DS astroglia exhibited significantly more (7–34-fold) Ca^2+^ transients than control isogenic astroglia (averaged number of calcium transients in a 5-min imaging session: DS1A: 58±6, DS4A: 275±34, DS2UA: 8±2, mean±s.e.m.; *P*<0.0001, unpaired t-test, *n*=9 imaging sessions) (Fig. 2b). The average amplitude (ΔF/F; DS1A: 1.45±0.2, DS4A: 0.98±0.15; *P*<0.01) (Fig. 2c) and frequency (transients/min; DS1A: 0.41±0.10, DS4A: 0.88±0.16; *P*<0.01) (Fig. 2d) of Ca^2+^ transients were significantly different between DS1A and DS4A, whereas the kinetics were similar (T_1/2_, s; DS1A: 8.59±1.01, DS4A: 6.98±0.90; *P*=0.18) (Fig. 2e). These disparities are potentially due to epigenetic changes between the cell lines.

**Figure 2.**
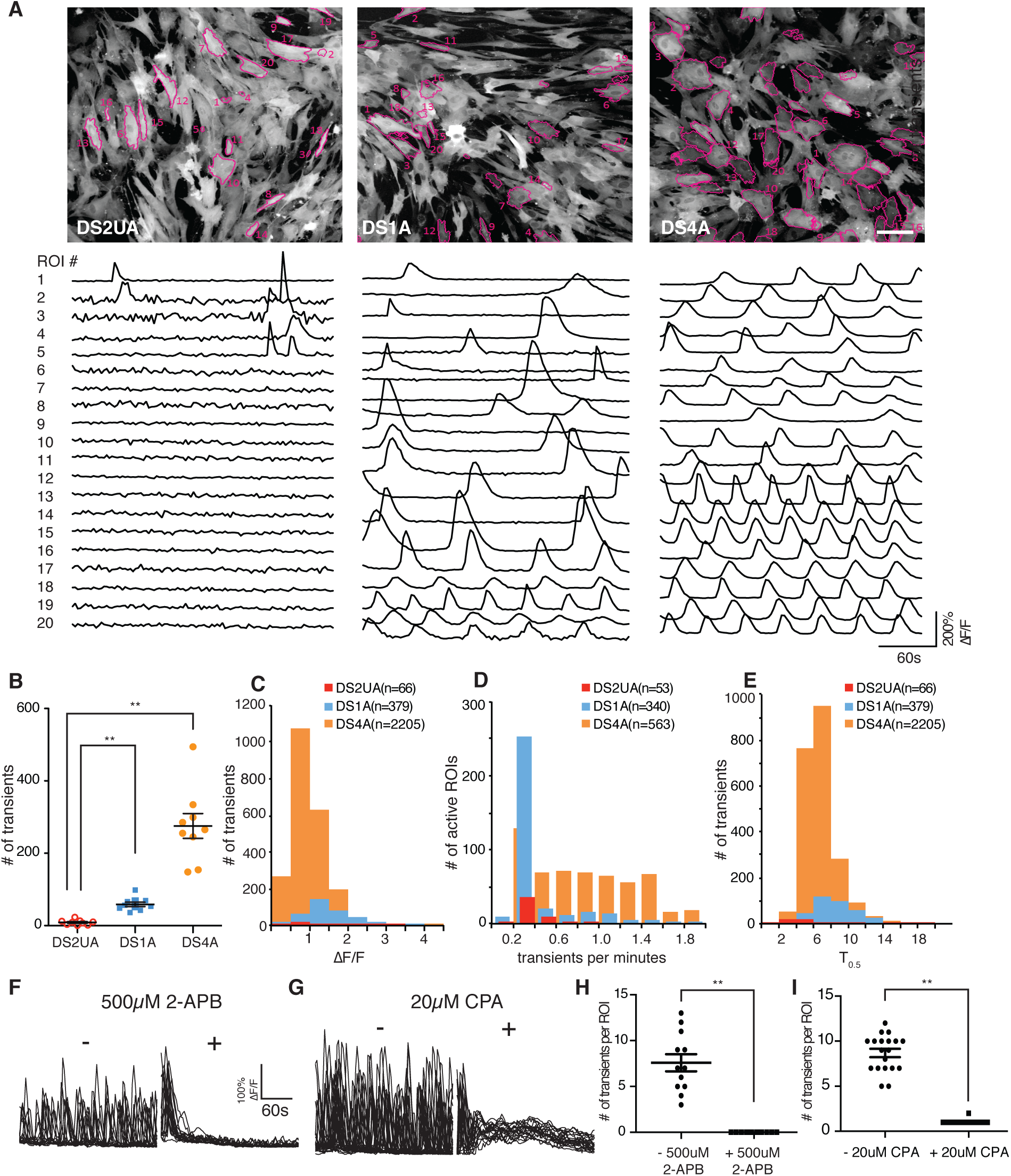
Imaging Ca^2+^ events in human iPSC-derived isogenic and DS astroglia. (a) Spontaneous Ca^2+^ responses in isogenic DS2UA and two DS astroglia (DS1A and DS4A). Representative ROIs (*n*=20) in the field of view showing Ca^2+^ fluctuations in DS2UA, DS1A, and DS4A. All ROIs were detected using FASP and marked with magenta outlines. Scale bars: 100 μm. (b) DS1A and DS4A displayed a significantly increased number of Ca^2+^ fluctuations in 5 min of imaging sessions, compared with DS2UA (9 independent imaging sessions). Features of Ca^2+^ fluctuations in DS astroglia (c–e): averaged kinetics (c), frequency (d), and propagation speed (e) of DS astroglia. Data were collected from 81 cells of DS1A and 188 cells of DS4A. (f-i) The Ca^2+^ fluctuations in DS4A could be abolished by incubation with IP_3_R antagonist (500 μM 2-APB, 17 ROIs, f & h) or depleting ER Ca^2+^ store (20 μM CPA, 23 ROIs, g & i) *P*<0.01 (**), unpaired t-test. Error bars represent mean±s.e.m.

Inositol triphosphate (IP_3_)-triggered Ca^2+^ release from the endoplasmic reticulum (ER) is considered a primary mechanism responsible for intracellular global Ca^2+^ waves^41^. Application of the IP_3_ receptor (IP_3_R) antagonist 2-aminoethoxydiphenyl borate (2-APB) abolished spontaneous Ca^2+^ fluctuations (Fig. 2f–i), as did depletion of intracellular stores by cyclopiazonic acid (CPA), suggesting that IP_3_-ER Ca^2+^ underlies both spontaneous and evoked events in DS astroglia.

Wavefront analysis of the spontaneous events revealed 33 clusters of cells (**Supplementary Fig. 3a**, left) in one field of view with temporally correlated Ca^2+^ fluctuations (**Supplementary Fig. 3a**, right). Cells within a waveform cluster were spatially intermingled, with identical distance distributions between temporally correlated and non-correlated cells (**Supplementary Fig. 3b**), suggesting that Ca^2+^ fluctuations do not propagate to adjacent cells. To further examine whether spontaneous fluctuations travel between cells, we performed Ca^2+^ imaging in a mixed culture of GCaMP6m-expressing control isogenic astroglia with unlabeled DS4A, in a variety of ratios. Culturing with DS astroglia did not significantly increase the number of Ca^2+^ transients in control isogenic astroglia, even with a 10-fold excess of DS4A (**Supplementary Fig. 3c**), suggesting that spontaneous Ca^2+^ fluctuations were not induced in previously silent control isogenic cells. In addition, application of 10 μM *n*-octanol, a gap junction blocker, showed no effect on Ca^2+^ fluctuations (**Supplementary Fig. 3d**). Taken together, these results indicate that the abnormal spontaneous Ca^2+^ fluctuations observed in DS astroglia are likely the result of cell-autonomous changes.

Previous studies reported that acutely purified human astrocytes acquire sensitivity to extracellular cues such as neurotransmitter ATP and glutamate^42^. To exclude the possibility that differences in functional maturation of differentiated astroglia contributing to suppressed neuronal excitability, we examined transmitter-evoked Ca^2+^ responses of DS astroglia and compared with isogenic controls. Both DS and control isogenic astroglia responded robustly to ATP (representative traces shown in **Supplementary Fig. 4a-b**) in terms of the number and amplitude of intracellular Ca^2+^ transients. Similarly, both DS astroglia and control isogenic astroglia responded to glutamate at micromolar concentrations (**Supplementary Fig. 4c-d**). Thus, DS and control astroglia respond similarly to neurotransmitters, further suggesting that Ts21 does not influence functionally maturation of differentiated astrocytes.

### Blocking spontaneous Ca^2+^ fluctuations in DS astroglia rescues suppressed neuronal excitability

We next tested whether the suppression of neuronal activity might be caused by the abnormally frequent spontaneous Ca^2+^ fluctuations observed in DS astroglia. Since pharmacological block of IP_3_ receptors abrogated spontaneous Ca^2+^ waves (Fig.2f-i), we knocked down (KD) the expression of *IP*_*3*_*R2*, the main *IP*_*3*_*R* isoform in astrocytes, with short hairpin RNAs (shRNAs) in DS astroglia DS4A. *IP*_*3*_*R2* KD, corresponding to ~50% knockdown (Fig. 3b), significantly reduced the number of active ROIs showing spontaneous Ca^2+^ transients [scrambled shRNA: 61.0±3.8; *IP*_*3*_*R2* shRNA-1: 21.3±.2.4 (35%), *IP*_*3*_*R2* shRNA-2: 14.3±1.8 (24%); *P*<0.001] (Fig. 3a,c), supporting the pharmacological results.

We next imaged the activity of neurons co-cultured with DS4A astroglia with knocked-down *IP*_*3*_*R2*. This rescued the reduced amplitude of evoked neuronal Ca^2+^ transients (measured as normalized ΔF/F; *IP*_*3*_*R2* shRNA-1: 0.91±0.0.4, shRNA-2: 0.93±0.03) to the level of isogenic control astroglia (1.01±0.04, *P*=0.28). In contrast, DS4A with no shRNA (0.60±0.05, *P*=0.0031) or control-scrambled shRNA (0.62±0.04, *P*=0.0018) showed significantly decreased neural activity (Fig. 3d). Therefore, elevated intracellular Ca^2+^ fluctuation mediated by IP_3_R2 is necessary to suppress neuronal excitability.

**Figure 3.**
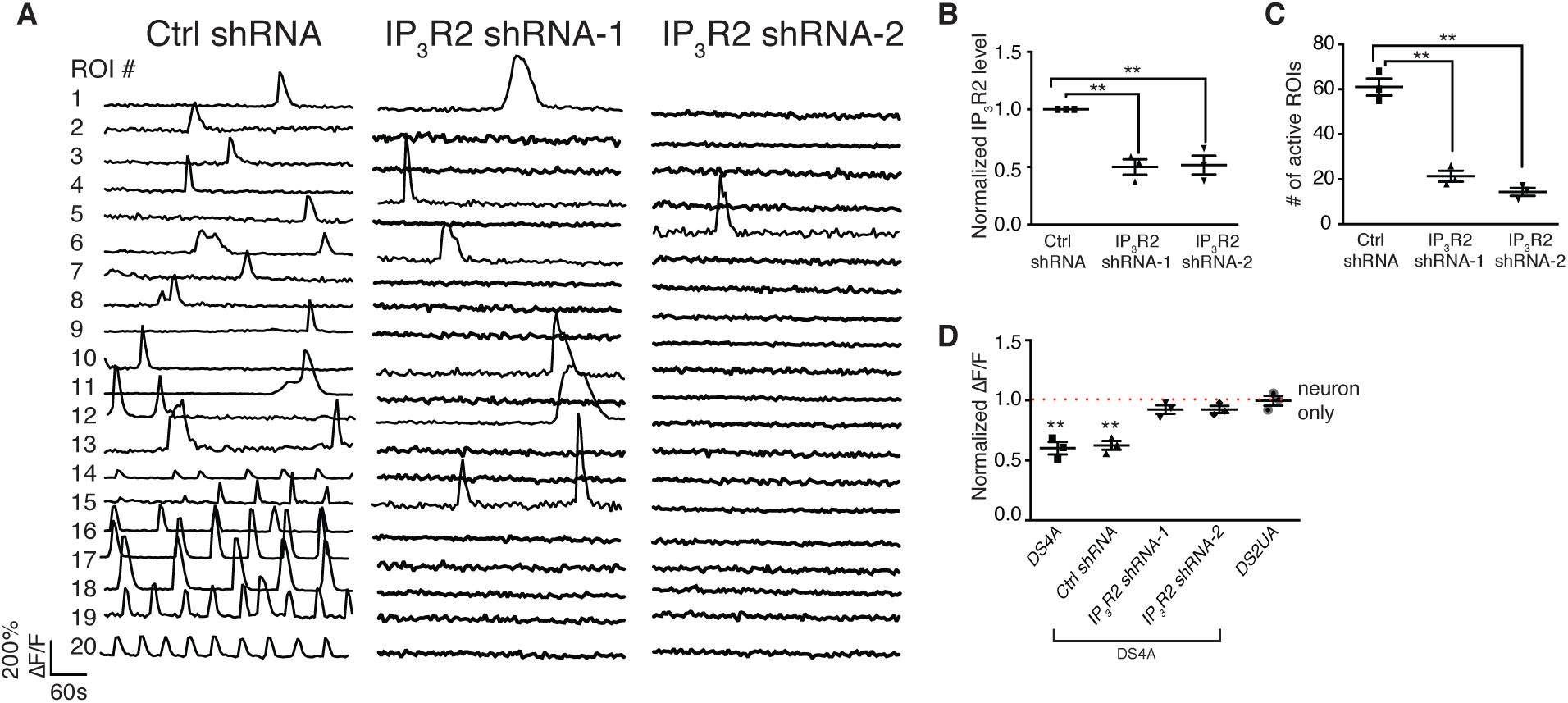
DS astroglial Ca^2+^ fluctuations are regulated by IP_3_R-ER pathway. (a–c) The Ca^2+^ events in DS4A were significantly decreased by knocking down the expression of IP_3_R. Representative ROIs (*n*=20) showing Ca^2+^ fluctuations in DS4A expressing scrambled shRNA (ctrl shRNA) and two shRNAs for IP_3_R (IP_3_R shRNA-1/2) (a). Real-time PCR confirmed the decreased expression of IP_3_R in the presence of IP_3_R shRNAs (3 RNA samples, b), corresponding to a decreased number of Ca^2+^ events in 5 min (3 imaging sessions, c). (d) Normalized fluorescence changes of H9 hESC-derived neurons in response to 40 FPs co-cultured with DS4A or DS4A expressing scrambled or IP_3_R shRNAs to those of neurons alone (dotted red line).

### Spontaneous Ca^2+^ fluctuations in DS astroglia are not driven by extracellular cues

As elevated spontaneous astroglia Ca^2+^ activity directly contributed to suppressed neuronal activity, we next sought to determine the factors driving elevated Ca^2+^ activity in DS astroglia. We first performed single-cell analysis of gene expression related to Ca^2+^ signaling pathways (mGluRs, purinergic receptors, GPCRs, and Ca^2+^ pumps; **Supplementary Fig. 5a**) in DS astroglia. We also monitored the expression of a panel of astrocytic markers to account for the differentiation state of individual cells (**Supplementary Fig. 5a-b)**. We then performed unsupervised clustering analysis of the cells by their gene expression patterns. We found that DS astroglia (e.g. DS4A) clustered into two groups (**Supplementary Fig. 5d**), distinguished by elevated expression of Ca^2+^ handling genes such as *ATP2B1*, *NCX1*, *RYR1/3*, *STIM1*, *NCLX*, *IP3R3*, *ORAI1*, and chromosome 21 gene *S100β* (**Supplementary Table 1**). This suggests that a subset of DS astroglia may display elevated spontaneous Ca^2+^ fluctuations. In DS astroglia, astrocytic markers such as *CD44*, *CX43*, *AQP4, NF1A*, and *ALDOC*, were not differentially expressed between the two clusters.

We next performed a similar analysis of gene expression patterns in control isogenic astroglia (e.g. DS2UA). In contrast, we failed to identify significant clustering (**Supplementary Fig. 5c**) of genes related to the Ca^2+^-handling toolkit.

Moreover, from the single-cell gene analysis, we found that metabotropic glutamate receptors (GRM1/2/3/4/5/6/7/8) and purinergic receptors were elevated in a subset of DS4UA. We next investigated whether spontaneous fluctuations in DS astroglia could be modulated by pharmacological manipulation of these receptors. ATP treatment led to a 2-fold increase in the frequency and a 1.4-fold increase in the amplitude of spontaneous Ca^2+^ fluctuations in ~40% of regions of interest (ROIs) (Fig. 4a). However, treatment with P2 isotype-specific ATP receptor antagonists (PPADS for P2X, MRS2179 for P2Y; Fig. 4b, **Supplementary Fig. 6b**), non-specific P2 antagonists (suramin; Fig. 4c), or an adenosine A_1_-receptor antagonist (DPCPX; Fig. 4d) had no significant effect on spontaneous Ca^2+^ fluctuations, suggesting that while ATP can modulate spontaneous Ca^2+^ events in DS astroglia, it is not sufficient to evoke them. CHPG (a selective mGluR5 agonist) showed no significant effects on amplitude, frequency, or kinetics of spontaneous Ca^2+^ fluctuations (**Supplementary Fig. 6a**). Similarly, mGluR5-selective (MPEP), non-selective mGluR (MCPG), and mGluR2/3-selective (LY341495) antagonists, as well as a glutamate transporter inhibitor (TFB-TBOA), also had no effect (Fig. 4e, **Supplementary Fig. 6c–e**). The TRPA1 channel antagonist HC030031 also had no significant effect on spontaneous Ca^2+^ fluctuations (Fig. 4f), consistent with the lack of microdomain Ca^2+^ activity observed^43^. In summary, while both intrinsically and extrinsically driven calcium transients depend on IP_3_-mediated release from ER stores, our results suggest that spontaneous fluctuations are unlikely to be driven, though can be modified, by extracellular cues.

**Figure 4.**
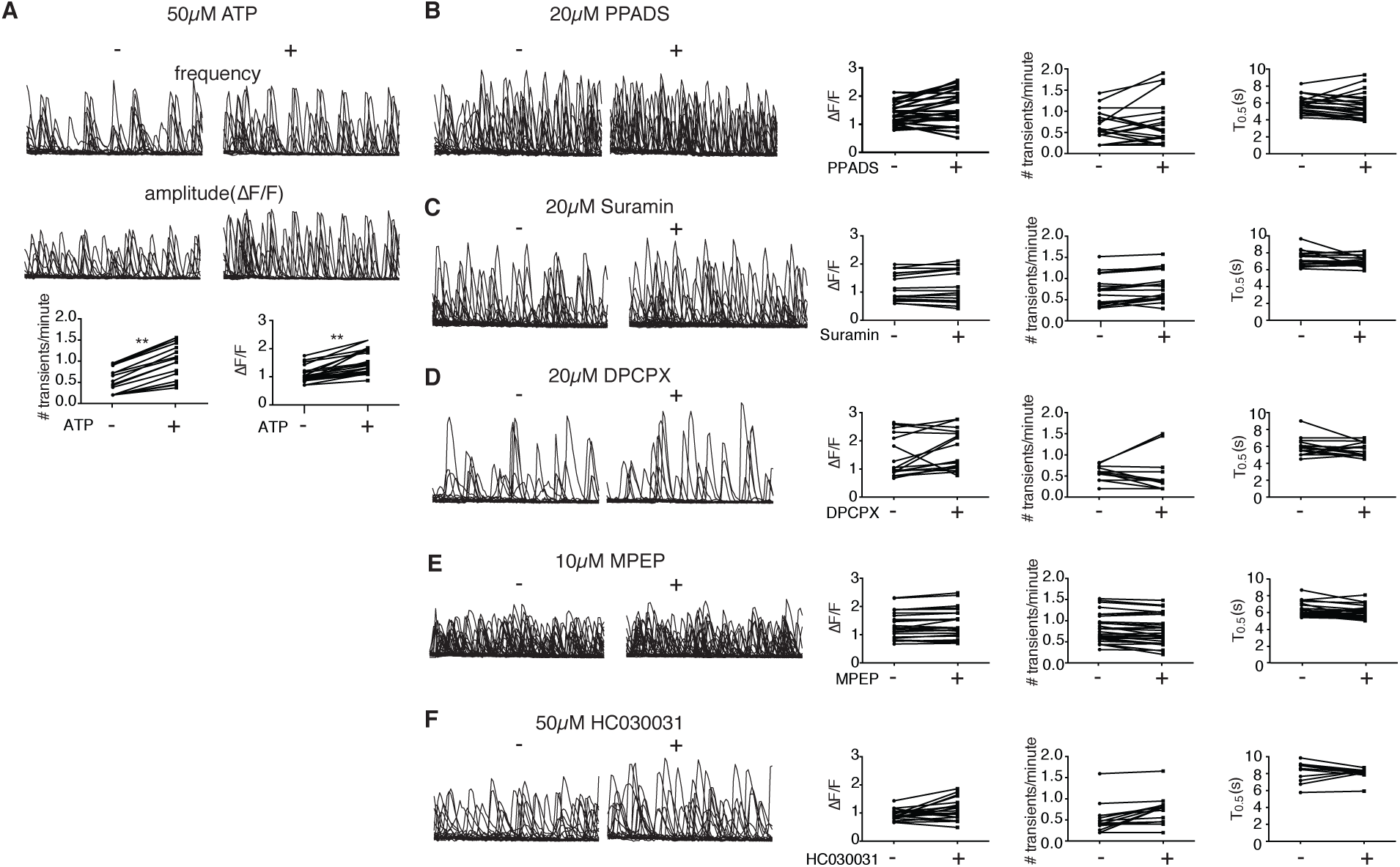
Spontaneous fluctuations in DS astroglia could not be modulated by pharmacological manipulation. (a) ATP increased the frequency and amplitude of Ca^2+^ events in previously active cells (20 ROIs were separately selected for either frequency or amplitude, *P*<0.01). (b–c) Purinergic receptor antagonist (20 μM PPADS, 25 ROIs, b), (20 μM suramin, 18 ROIs, c) failed to modulate the Ca^2+^ fluctuations in DS4A. (d) A1 adenosine receptor antagonist (DPCPX 20 μM, 18 ROIs), (e) mGluR5 antagonist (10 μM MPEP, 42 ROIs), and (f) TRPA inhibitor (50 μM HC030031, 20 ROIs) failed to modulate the Ca^2+^ fluctuations in DS4A. Left: representative traces. Right: quantification of amplitude, frequency, and kinetics. *P*<0.01 (**), unpaired t-test. Error bars represent mean±s.e.m.

### S100β regulates spontaneous Ca^2+^ fluctuations in DS astroglia

From gene analysis, we also noticed that S100β, a Ca^2+^-binding protein located on HSA21 and primarily expressed in astrocytes, was one of the top genes differentially expressed across DS4A cells. Previous studies reported that astrocytes release S100β, eliciting neurotrophic effects and regulating synaptic plasticity and rhythmic neuronal activity by chelating extracellular calcium^44,45^. Given the particular interest in the context of Ts21 DS, we thus asked the question whether elevated S100β might contribute to the spontaneous intracellular Ca^2+^ fluctuations in DS astroglia.

We first quantified the expression level of S100β in Ts21-derived astroglia. qPCR analysis showed an averaged 11-fold greater expression of *S100β* in DS astroglia (DS1A and DS4A) compared with control isogenic DS2UA cells (**Supplementary Fig. 1c**). Expression of S100β protein was enriched in DS astroglia compared to DS2UA (Fig. 5a–b; 9.9- and 10.7-fold increased expression S100β, for DS1A and DS4A, respectively, compared to DS2UA).

We next selectively knocked down *S100β* in DS4A, and performed Ca^2+^ imaging. We co-expressed mCherry as a proxy for the extent of *S100β* KD and used fluorescence-activated cell sorting (FACS) to select the top 15% of cells with potent *S100β* KD and use the bottom 15% of cells as a control group with normal *S100β* levels (**Supplementary Fig. 7a**). The *S100β* KD population contained ~10-fold lower *S100β* levels compared to the control group (*P*<0.001) (Fig. 5d), indicative of effective *S100β* KD. *S100β* KD led to a 3.5-fold decrease in spontaneous Ca^2+^ transients during a 5-minute window (*P*<0.001; Fig. 5c, e). These data suggest that S100β modulates spontaneous Ca^2+^ fluctuations in DS astroglia.

**Figure 5.**
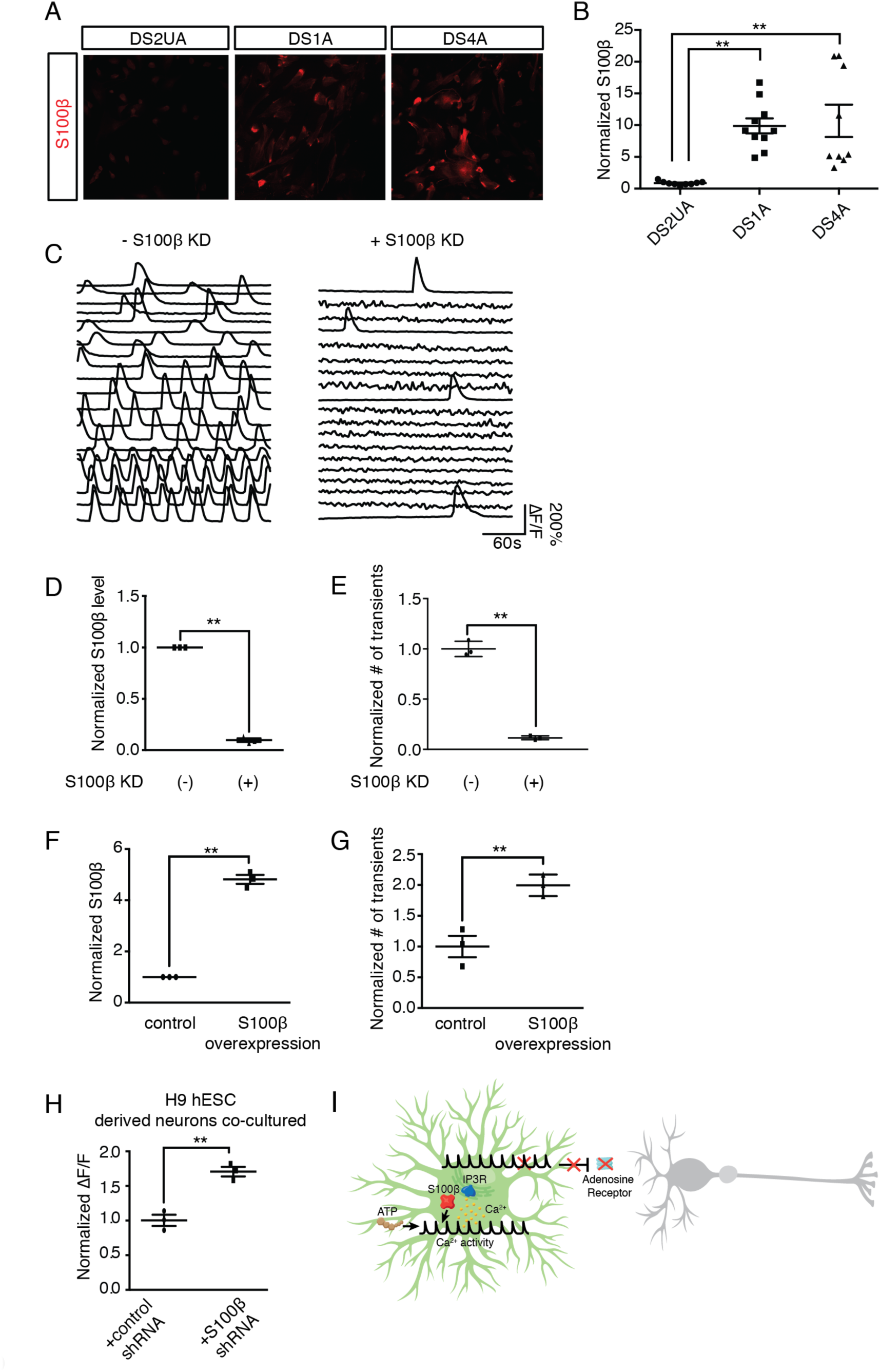
S100β regulates spontaneous Ca^2+^ fluctuations in DS astroglia. (a-b) Immunostaining of S100β in iPSC-derived astroglia revealed increased expression in DS astroglia compared with isogenic DS2UA (3 images of immunostaining, 12.5±1.0% for DS2UA, 80.4±1.8% for DS1A, 75.3±2.9% for DS4A, *P*<0.01). Scale bar: 50 μm. (c) Representative ROIs (*n*=20) showing spontaneous Ca^2+^ fluctuations in populations of DS4A with and without S100β KD. Scale bars: 50 μm. (d-e) The number of Ca^2+^ events was significantly decreased in DS4A populations with decreased expression of S100β. The normalized S100β expression levels (3 RNA samples, d) and the number of Ca^2+^ events (3 imaging sessions of 5 min, e) with and without S100β KD are shown. (f–g) Overexpression of S100β increased the number of Ca^2+^ events in DS1A. qPCR analysis confirmed elevated expression of S100β in DS1A when S100β was overexpressed (3 RNA samples, 4.8±0.2–fold relative to empty vector group, *P*<0.01, f). 2-fold more Ca^2+^ events in 5 min were detected in DS1A when S100β was overexpressed (3 imaging sessions, g). (h) Blocking intracellular Ca^2+^ events by S100β KD increased activity of H9 hESC-derived neurons co-cultured with DS4A. The fluorescence changes of H9 hESC-derived neurons in response to 40 FPs stimuli co-cultured with 2 populations of DS4A are shown. (i) Hypothetic model. DS astroglia exhibit aberrant Ca^2+^ fluctuations, which are dependent on IP_3_R-ER pathway, and can be modulated by ATP and HSA21 gene *S100β*. The elevated Ca^2+^ fluctuations inhibit neuronal activity, which can be rescued by blocking either aberrant Ca^2+^ fluctuations or adenosine receptors. *P*<0.01 (**) or 0.05 (*), unpaired t-test. Error bars represent mean±s.e.m.

Given the reported role of secreted S100β protein in modulating neural activity, we incubated the cultures with antibodies against S100β or Tuj1 (without permeabilization). After 10 minutes incubation, there was no effect on spontaneous Ca^2+^ events of either antibody (**Supplementary Fig. 7b**), suggesting that the spontaneous Ca^2+^ events are mediated by intracellular S100β.

We then asked whether overexpression of S100β protein would also modulate spontaneous Ca^2+^ fluctuations. We overexpressed *S100β* in DS1A (Fig. 5f), in which the number of spontaneous Ca^2+^ transients is less abundant than DS4A. After two days of expression, we observed a 2-fold increase in the number of Ca^2+^ transients (*P*<0.01; Fig. 5g). Thus, we conclude that increased cytosolic S100β expression is both necessary and sufficient to drive the spontaneous Ca^2+^ fluctuations observed in DS astroglia.

Finally, we examined whether DS astroglia with spontaneous Ca^2+^ fluctuations alleviated by *S100β* KD still suppressed neuronal excitability. We recorded evoked Ca^2+^ events in response to FP stimuli in H9 neurons co-cultured with DS4A with or without *S100β* KD. H9 neurons co-cultured with DS4A with potent *S100β* KD displayed significantly larger (1.7-fold; *P*=0.0027) neural activity than those without *S100β* KD (Fig. 5h), suggesting that *S100β* KD successfully rescued neuronal activity suppressed by DS4A. Thus, together with the findings above (Fig. 3d), we conclude that blocking Ca^2+^ fluctuations in DS astroglia by genetic ablation of either *IP*_*3*_*R2* or *S100β* is sufficient to rescue the excitability decreases of co-cultured neurons.

In summary, our data indicate the functional importance of astrocyte-neuron interplay in regulating neuronal excitability in a DS-iPSCs based model. The aberrant Ca^2+^ fluctuations in human DS astrocytes depend on intracellular IP_3_-ER Ca^2+^ release and are mediated by the overexpression of HSA21 protein S100β. Blocking spontaneous Ca^2+^ fluctuations in DS astroglia or adenosine-mediated astrocyte-neuron signaling successfully rescued suppressed neuronal excitability (Fig. 5i).

## Discussion

Combining human stem cell technology with genetically encoded Ca^2+^ indicators and quantitative analysis tools provides a powerful platform to study neuron-astrocyte interaction, in both physiological and pathological conditions, especially at early developmental stages. Using this platform, we imaged and characterized the effect of Ts21-iPSC–derived astroglia on neuronal networks. DS astroglia produced structural and functional deficits in co-cultured neurons. Specifically, neurons co-cultured with DS astroglia displayed decreased global excitability. Such decreased global excitability of neurons corresponded with increased amplitudes of post-synaptic activity and synaptic density, consistent with accepted mechanisms of homeostatic synaptic plasticity and synaptic scaling^46^. More importantly, our data is in line with a rodent DS model (overexpression of Ts21 gene, *Dyrk1a*) study, in which the dendritic spine density and mEPSC amplitude increased while frequency of mEPSCs remained unchanged in prefrontal cortical pyramidal neurons.^30^ It is worth investigating whether this alteration of synaptic properties could also be imposed by astrocytes in the Dyrk1a rodent model.

We further showed functional differences between DS astroglia and control isogenic astroglia in terms of intracellular Ca^2+^ dynamics. We observed elevated spontaneous Ca^2+^ fluctuations that are frequent and periodic only in DS-derived astroglia, but not in an isogenic control cells. These aberrant Ca^2+^ fluctuations in DS astroglia are necessary to drive suppression of global excitability in co-cultured neurons, as evidenced by rescue by genetic or pharmacological block.

What causes aberrant Ca^2+^ fluctuations in DS astroglia? In the present study, we demonstrate that overexpression of cellular S100β in DS astroglia mediates elevated spontaneous Ca^2+^ fluctuations, which subsequently regulate neuronal excitability (Fig. 5g-h). This finding is of particular interest, as S100β is a Ca^2+^-binding protein. Previous research^47^ has shown that secreted S100β stimulates a rise in intracellular Ca^2+^ concentration in both neurons and glia. Furthermore, extracellular S100β regulates the firing patterns of neurons by reducing extracellular Ca^2+^ concentrations^44^. In our studies, extracellular S100β did not influence spontaneous Ca^2+^ fluctuations in DS astroglia, whereas cytosolic of S100β did. Further investigation is necessary to parse the various functions of secreted and cytosolic S100β in healthy and disease-model astrocytes and neurons.

A major open question in DS research is the mechanism by which the overdose of hundreds of genes on HSA21 disrupts brain function. To date, several candidate genes have been identified, including *DYRK1A*, *SIM2*, *DSCAM*, *KCNJ6*, *NKCC1*, and *miR-155*^1,48,49^ (**Supplementary table 1**). Overexpression of S100β, at the distal end of the HSA21 long arm, has been shown to generate reactive oxygen species (ROS)^25^ in hiPSC-derived DS astroglia, leading to neuronal apoptosis^3^. Previous research reported that ROS induce lipid peroxidation, activate the PLC-IP_3_R pathway, and cause Ca^2+^ increases in astrocytes^50^. Indeed, we found that spontaneous Ca^2+^ activity was mediated by IP_3_R2-regulated ER stores. Though we do not have direct evidence to link S100β, ROS, and PLC-IP_3_R, S100β might mediate perturbed Ca^2+^ dynamics via ROS in DS astroglia.

Our study provides additional evidence to support the hypothesis that astrocytic Ca^2+^ signaling modulates neural activity, critical for brain function during development. A grand challenge is to elucidate the pathways regulating astrocyte-neuron interplay during development. In the present study, our results indicate that astrocyte-neuron interaction via purinergic signaling might be a significant contributor linking aberrant astrocytic Ca^2+^ to neuronal functional deficits in DS. We showed that treatment with DPCPX, an adenosine A_1_ receptor antagonist, rescued the suppressed Ca^2+^ activity of H9 hESC-derived neurons co-cultured with DS astroglia (Fig. 1i). To what extent ATP potentiates and/or inhibits neuronal activity is still under debate; however, adenosine predominantly inhibits synaptic activity via A_1_ receptors^33–35^.

In conclusion, the combination of a human iPSC DS model with functional imaging, and pharmacological and genetic manipulation provides a platform for quantitative measurement of human cellular physiology and for mechanistic studies of disease pathophysiology. Though animal models of neurological disorders play an important role in studying the effects of specific genetic and experimental perturbations and in testing potential treatments, they often fail to faithfully recapitulate the full spectrum of human phenotypes, which can lead to false conclusions owing to molecular and cellular differences between the systems. Future improvements to iPSC models will include 3-dimensional culture^51^, multi-color imaging, and incorporating genetically encoded indicators for other molecules and cellular states (*e.g.* glutamate)^52^. Our imaging platform can be applied to the study of other neurological diseases, as well, even to the level of testing specific drug combinations on neuron-astrocyte co-cultures developed from single healthy or diseased individuals.

## Materials and Methods

### Plasmid construction

*IP*_*3*_*R2*, *S100β*, and scrambled shRNA KD plasmids were ordered from Sigma (MISSION® shRNA Library, pLKO.1 with *U6* promoter driving shRNA expression). Lentiviruses were produced in HEK293T cells and used to infect astrocytes. To construct shRNA-mCherry plasmids, shRNA plasmids were digested with KpnI and BamHI (New England BioLabs; Ipswich, MA). mCherry flanked by KpnI and BamHI was ligated into the shRNA vector. To construct *PGK* promoter-driven S100β, *S100β* was amplified by PCR using astrocytic cDNA as a template, digested with KpnI and BamHI, and then ligated to plasmids digested with KpnI and BamHI.

### Neural differentiation of human ESCs and iPSCs

H9 human ESCs were obtained from WiCell (Madison, WI). Control isogenic trisomy 21 and euploid iPSCs, DS1, DS2U, and DS4, were engineered in Dr. Anita Bhattacharyya’s lab, as previously described^4^. H9 ESCs and iPSCs were maintained on matrigel (Becton-Dickinson, 356234) in mTeSR1 medium (StemCell Technologies, 05850). Mycoplasma contamination was tested for routinely. We used previously described protocols for neural differentiation^29^, with minor modifications. Inhibitors of SMAD signaling (10μM SB431542 and 100 nM LDN193189, both from Tocris) were added for the first 6 days to promote neural induction^28^.

### Derivation and culture of astrocytes

Control isogenic and DS iPSCs were differentiated into neural progenitors and cultured as spheres for 3 months. The astrospheres were attached to fibronectin-coated dishes (Sigma, F0895), dissociated into single cells, and cultured in an optimized commercial medium for human primary astrocytes (ScienCell Research Laboratories, 1801). Human primary astrocytes (HA) were also from ScienCell Research Laboratories (1800). We performed karyotype analysis to confirm the trisomy states of DS1- and DS4-derived astroglia (DS1A and DS4A), and the disomy state of control isogenic line DS2U-derived astroglia (DS2UA), prior to and after the Ca^2+^ experiments using a service provided by Cell Line Genetics. Indeed, chromosome alteration usually occurs more frequently during the maintenance of iPSCs before differentiation^53^. The cell size was analyzed by randomly selecting 5 cells from 3 bright field images. Pixel areas of each selected cell were calculated and averaged in ImageJ.

### Lentivirus production

Lentiviruses were produced by co-transfecting HEK293T cells (ATCC) with 5 μg pSIV-*Synapsin-1*-GCaMP6m or pHIV-*EF1α*-Lck-GCaMP6m, scrambled or S100β shRNAs, 2 μg pHCMV-G, and 3 μg pCMV-deltaR8.2, using 40 μl SuperFect (Qiagen, 301305). Supernatant containing viral particles was collected, filtered, and concentrated 72 h later with an Ultra-4 centrifugal filter (Millipore, UFC810024).

### Ca^2+^ imaging and analysis in astrocytes

Primary astrocytes or iPSC-derived astrocytes were seeded onto 8-well slides (Ibidi, 80826, optically clear), coated with fibronectin and infected with lentiviruses encoding GCaMP6m driven by the *EF1α* promoter, then subjected to Ca^2+^ imaging. For *IP*_*3*_*R2* KD, DS4A cells were infected with lentiviruses encoding shRNA and GCaMP6m; Ca^2+^ imaging followed. For *S100β* KD, DS4A cells were infected with lentiviruses encoding shRNA, sorted into 2 populations by FACS according to mCherry intensity, and infected with GCaMP6m for each population; Ca^2+^ imaging followed. For each cell line, 3 Ca^2+^-imaging sessions (each session contains 3 fields of view) were collected from independent samples. For mixed cultures of control isogenic and DS astrocytes, control isogenic DS2UA were first infected with lentiviruses expressing *EF1α*- GCaMP6m, then seeded with DS4A, followed by Ca^2+^ imaging. Three days post-infection, frame scans were acquired at 2 Hz (512x512 pixels) for a period of 300 s using a Zeiss LSM 710 confocal microscope (× 20 magnification, N.A.=0.8 objective). Agonists or antagonists (Tocris) were added at frame 10 during continuous imaging. For quantification of ATP and glutamate-evoked activity, to eliminate the confound of spontaneous activity, only ROIs that were silent during the initial imaging period were analyzed for a response to added ATP or glutamate. Furthermore, we ensured that these evoked responses were time-locked to agonist application.

Because of these complex spatiotemporal patterns of Ca^2+^ dynamics in astrocytes, we developed a computational tool, named FASP^40^, to quantitatively and automatically analyze the large-scale imaging datasets to ensure that the analysis is identical and objective for all cells and across experiments. As an unsupervised analytic method, FASP is data-driven, learning model parameters using machine-learning techniques to automatically detect ROIs displaying Ca^2+^ fluctuation. In addition, designed under probabilistic principles, FASP has strong statistical power to detect weak signals (ROIs) that are easily ignored by purely manual analysis. Our simulation study verified that some ROIs with weak signals were ignored by manual analysis but correctly detected by FASP. By judicious application of various statistical theories, FASP confers tuning parameters with probabilistic meaning, which can be directly translated into the false discovery rates. This algorithm greatly facilitates the usability of parameter settings and ensures the reproducibility of the results and equal comparison across experiments.

Specifically, we set a single threshold corresponding to a false discovery rate of 0.01; that is, an average of 1% of all identified active ROIs are expected to be false positives. The threshold is fixed for all experiments and conditions.

Given a time-lapse astrocytic Ca^2+^-imaging data set, FASP generates a set of ROIs and corresponding characteristic curves. For each pixel in an ROI, there is a corresponding activity curve for which the time shift with respect to the characteristic curve is also estimated. Based on the results of FASP, we quantified various parameters of astrocytic Ca^2+^ signals according to the following:

- The signal-to-baseline ratio of fluorescence was calculated as 

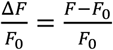

where the baseline fluorescence *F*_0_ is estimated as the 10^th^ percentile of the fluorescence level over all time points of the measurement.
- The number of Ca^2+^ transients is calculated as the number of peak responses from all ROIs detected in each time-lapse imaging session.
- The number of active ROIs is calculated as the total number of ROIs detected by FASP in the field of view of each time-lapse imaging session.

#### Amplitude

To calculate the amplitude of a Ca^2+^ transient we first transformed the raw time-intensity curves into signal-to-baseline ratio of fluorescence (ΔF/F0=(F-F0)/F0), where the baseline fluorescence F0 is estimated as the 10^th^ percentile of the fluorescence levels (intensities) at all the time points during measurement.

#### Frequency

To calculate the frequency of Ca^2+^ fluctuations more reliably, we first determined the average duration between 2 contiguous events, and then defined the frequency as the inverse of the average duration. For those ROIs that only displayed single Ca^2+^ transients during the imaging session, the information contained in the single-event time series is insufficient for point estimation of frequency. These ROIs are expected to have a positive frequency between 0 and 0.2 transients per minute.

#### T_0.5_

Decay kinetics or T_0.5_(off) was calculated using linear interpolation as the time from peak to half-amplitude of an event.

#### Propagation speed wavefront analysis

On the basis of estimated pixel-wise time shifts from the characteristic curve, wavefronts of Ca^2+^ transients were located; accordingly, the propagation speed of Ca^2+^ events within an ROI was obtained by estimating the average distance between wavefronts. Active ROIs detected in DS4A were divided into 33 clusters of timed coincidence by unsupervised clustering analysis (Affinity Propagation Clustering Algorithm)^54^. Any pairs of ROIs within the same cluster were recognized as highly coincident, while any pairs of ROIs from 2 different clusters were recognized as weakly coincident. Distributions of pixel distance of correlated and uncorrelated pairs were then measured and plotted.

### Neuron-astrocyte co-culture and imaging

Neurospheres were seeded on matrigel-coated glass-bottom dishes (MatTek, P35G-1.0-14-C), and cultured in neuronal medium [neurobasal medium, 21103-049; 1% N-2 supplement, 17502-048, 2% B-27 supplement, 17504-044; 10 ng/ml BDNF (450-02) and GDNF (450-10)] for 40 days. The medium components were purchased from Thermo Fisher Scientific, and cytokines from Peprotech. Neurons were then infected with lentiviruses expressing *Synapsin-1*-GCaMP6m. Two days post-infection, astrocytes were seeded on top of neurons to establish co-culture. After 3–7 days, infected neurons were stimulated using a custom-built field stimulator with platinum wires and imaged using a Zeiss LSM 710 confocal microscope (× 20 magnification, 0.8 NA, 512×512 pixels, 458 ms/frame). Field stimuli were delivered as 40 V, 30 Hz, 1 ms pulses for the following trains: 10, 20, 40, 80 field stimuli in HBSS with 2mmol CaCl_2_ and MgCl_2_. When chemicals were used, they were applied 3 days prior to imaging, except DPCPX, which was acutely applied 1 h prior to imaging. All chemicals were purchased from Tocris. All neuronal imaging experiments were repeated 3 times, and 10 ROIs were selected for analysis using a customized script (FluoAnalyzer) in MATLAB (MathWorks). ROIs (*n*>10 for each imaging file) were manually selected, and the fluorescence intensity (F) at each frame was quantified as the mean of all selected ROIs. The neuronal responses were calculated as ΔF/F (F-F_0_/F_0_), where F was quantified as the mean of all selected ROIs (*n*>10 in each field of view), and F_0_ was taken as the mean of all ROIs across the first 3 frames.

### Immuncytochemistry

Cells maintained on cover glasses (Fisher Scientific, 12-545-81) were washed with PBS 3 times before being fixed with 4% paraformaldehyde (VWR, 100503-916) for 15 min. After washing, cells were treated with 0.1% Triton X-100 (Fisher Scientific, BP151-500) for 10 min, blocked with 10% bovine serum albumin (Sigma, A9647) for 60 min, and incubated with primary antibodies at 4 °C overnight followed by secondary antibodies for 1 h at room temperature. Cells were washed with PBS 2 times after each antibody incubation and mounted on glass slides (Fisher Scientific, 12-550-123) using ProLong^®^ Gold Anti-fade Mountant with DAPI (Thermo Fisher Scientific, P36935). Primary antibodies used included: AQP4 (Santa Cruz Biotech, sc-20812, rabbit), CD44 (Abcam, ab6124, mouse), GFAP (Millipore, MAB360, mouse; AB5840, rabbit), TUJ1 (COVANCE, MMS-435P, mouse), Synapsin-I (Millipore, AB1543, rabbit), S100β (Abcam, ab11178, mouse), and PSD95 (NeuroMab, K28/43, mouse). Secondary antibodies included Alexa488-conjugated donkey anti-rabbit (A21206) and Alexa594-conjugated goat anti-mouse (A11005), and were purchased from Thermo Fisher Scientific.

### Immunocytochemistry analysis

Images were obtained using a Zeiss LSM 710 confocal microscope (× 40 magnification, N.A. 1.3 oil objective). All immunostaining experiments were performed 3 times, and representative results were presented.

Puncta density quantification: Using the spot-detection feature in the Imaris software (Bitplane) the number of colocalized Synapsin-1 and PSD95 per μm of dendrite was obtained to calculate the puncta density. S100β immunocytochemistry analysis: Using FIJI the fluorescence intensity of each imaging field was analyzed.

### mEPSC recordings

Whole-cell voltage clamp experiments were performed 17–19 days after plating. mEPSCs were recorded in an external solution containing 140 mM NaCl, 5 mM KCl, 10 mM HEPES, 2 mM CaCl2, 1 mM MgSO4, 1 μM tetrodotoxin (TTX), 50 μM AP-5, and 20 μM bicuculline (pH 7.4 with NaOH, 290 mOsm/l). Borosilicate glass electrodes were filled with an internal solution containing 145 mM CsCl, 1 mM EGTA, 5 mM HEPES, 0.1 mM CaCl2, 2 mM MgSO4 (PH 7.4 with CsOH, 275 mOsm/l). The seal resistance was greater than 1 GΩ and the series resistance was no greater than 20 MΩ. All recordings were made with an Axopatch 200B patch-clamp amplifier (Axon Instruments, Foster City, CA, USA). Whole-cell currents were filtered at 2 kHz and digitized at 10 kHz. All neurons were voltage-clamped at –60 mV.

### mEPSC analysis

The mEPSC events were detected with Mini Analysis software (Synaptosoft Inc., Fort Lee, NJ, USA). The accuracy of detection was confirmed by visual inspection.

### RNA isolation and qPCR

Total RNA was prepared from cells *(n*=3) with RNeasy kit (Qiagen, 74104). Complementary DNA was prepared with iScript RT Supermix (Bio-Rad, 170-8841). qPCR was performed with iTaq™ Universal SYBR^®^ Green Supermix (Bio-Rad, 172-5121) on a CFX96™ Real-Time System (Bio-Rad), and the data was collected with Bio-Rad CFX Manager 3.0. Gene expression levels were quantified relative to the housekeeping gene, *GAPDH*.

### Single-cell expression analysis

DS astrocytes were digested and then sorted by FACS to get rid of cell debris and dead cells. The cell suspension was loaded onto a C1 Single-Cell Auto Prep Array for mRNA Seq (10–17 μm; Fluidigm, 100-5760), and single cells were captured and lysed to get cDNA on Fluidigm’s C1 platform. Gene expression patterns of single cells (*n*=46) were studied using the 48.48 Dynamic Array Chip for Gene Expression (Fluidigm, BMK-M48.48), which assembles cDNA from individual cells to create individual qPCR reactions following the manufacturer’s instructions.

The values of gene expression were pre-processed by taking the inverse, applying a square-root transformation, and rescaling the expression to zero mean and unit variance. The similarity matrix was computed first using the default method of negative distance (default parameters), and affinity propagation clustering was applied by setting the desired number of clusters to 2 in the R package, Apcluster.

The single-cell expression analysis consisted of 4 major components. First, pre-processing was conducted to impute missing values and make sure the expression values were approximated well by Gaussian distributions to facilitate follow-up analysis. Second, clustering analysis was done to discover groups within the cell populations. Third, significance tests were implemented to determine whether the resultant clusters were purely due to chance. Fourth, differential analysis was used to find the genes underpinning the clusters. Detailed discussion about these 4 components follows.

#### Preprocessing

In the raw data, the value for each gene denotes how many amplification cycles were required to cross the threshold, which is set using the AutoGlobal method. In our data, the maximum observed value was 29. A missing value indicated that the corresponding gene had too little expression to be amplified to reach the threshold quantity. In the raw data, missing values were marked by 999. We replaced all missing values by 60, which was around 2 times the maximum value observed. Then, the inverse of the values was used to represent the amount of expression. A square-root transformation was applied to each gene to normalize for expression-level differences. Finally, for each cell, all genes were normalized to have zero mean and unit variance to highlight differences between cells.

#### Clustering analysis

Affinity propagation clustering (APC) was applied. The algorithm was implemented in the R package, Apcluster. The algorithm requires users to input a similarity matrix. The default settings were adopted; in other words, Euclidian distance was calculated based on the data matrix and the negative distance was used as the similarity matrix. To be consistent with the observation that there were 2 groups of astrocytes, one with active Ca^2+^ fluctuations and the other one without, the desired number of clusters was set to 2. Notably, we did not know which cells were active, and the analysis was unsupervised.

#### Assessing statistical significance of resultant clusters

Since a clustering algorithm can always generate clusters even if there are no clusters actually present in the data, we sought to evaluate whether the resultant clusters were purely by chance. The null hypothesis was that there were no groups of cells that were closer within-group than between-groups (i.e., distances between cells were uniformly distributed). Permutation was used to generate the distribution for the null hypothesis. All genes in the data set were permuted 100,000 times, resulting in 100,000 data sets following the null hypothesis. For each resulting data set, we ran APC to get 2 groups. In APC, the objective function was the overall similarity. A histogram was obtained based on the 100,000 overall similarities. The position of the observed overall similarity indicated the significance of the observed value.

#### Differentially expressed genes between clusters

Standard differential analysis, such as t-test between groups, cannot be applied here because the clustering was based on all genes; hence, each gene was biased toward differential expression between clusters. To correct for this bias, a permutation procedure was designed. To test the significance for each gene, we shuffled the values in that gene 10,000 times while keeping all other genes fixed. In this way, the gene would not interfere with the clustering results, so there would be no bias. Each time we ran the clustering algorithm APC to get 2 groups. Based on the new clustering results, the significance of the gene was recorded and summarized into a histogram, which could be further used to derive the corrected *P* value. For example, if the original *P* value was 0.01, corresponding to the 2^nd^ percentile in the histogram, the corrected *P* value would be 0.02.

### Fluorescence-activated cell sorting (FACS)

DS astrocytes were infected with lentiviruses expressing S100β-shRNA-mCherry and collected 3 days later for sorting, which was performed by the FACS core at UC Davis. The top 15% of cells expressing high amounts of mCherry measured on fluorescence intensity were collected as mCherry “high”, and the bottom 18% of cells expressing low amounts of mCherry were collected as “low”. High mCherry fluorescence represents high expression of S100β shRNA and less expression of S100β.

### Inhibiting extracellular S100β

S100β (Abcam, ab11178, mouse) and TUJ1 (COVANCE, MMS-435P, mouse) antibodies (diluted 1:1000) were used to pretreat DS astrocytes. Following a 10 minute incubation cells infected with pHIV-*EF1α*-Lck-GCaMP6m were subjected to Ca^2+^ imaging.

### Karyotype analysis

Karyotype analysis was performed by Cell Line Genetics (Madison, WI).

### Statistical analysis

All values are shown as mean±s.e.m. To determine significant differences between groups, comparisons were made using a two-tailed unpaired t-test. For the modulation of Ca^2+^ fluctuation by ATP, two-tailed paired t-test was used. For mEPSC analysis, a one-way ANOVA was used to compare mEPSC amplitude and frequency among groups, followed by Fisher’s LSD pairwise comparison when appropriate. For single-cell expression analysis, a permutation test was applied for unsupervised clustering, and the differences of each gene between the two clusters were determined using two-sample unpaired Wilcoxon rank-sum test. A *P* value smaller than 0.05 was accepted for statistical significance. The sample size for each experiment was determined either by power analysis (2-Sample, 2-Sided Equality) or by referring to the sample size in similar studies^3,42^. For Ca^2+^ imaging experiments, imaging sessions were collected from at least 3 batches of cells, and ROIs were selected either automatically by FASP for astrocyte Ca^2+^ imaging or manually for neuronal Ca^2+^ imaging. For gene expression, RNA samples from three batches of cells were used. For immunostaining analysis, three batches of cells were fixed and five fields of view from each sample were selected for imaging. No randomization or blinding was used. No data was excluded.

## Acknowledgments

This work was supported by the Hartwell foundation (L.T.), NIH DP2 MH107059 (L.T.), and NIH R03 HD064880 (A.B.). This project was supported by the University of California, Davis, Flow Cytometry Shared Resource Laboratory; and with technical assistance from Ms. Bridget McLaughlin and Mr. Jonathan Van Dyke. We would like to give special thanks to Dr. Bart Borghuis for generously sharing the FluoAnalyzer codes, Dr. Karen Zito, Dr. Tommaso Patriarchi and Brian McGrew for their critical input and Lisa Makhoul for editorial assistance.

## Author Information

The authors declare no conflicts of interest.

